# Planning to revisit: neural activity in refixation precursors

**DOI:** 10.1101/660308

**Authors:** Andrey R. Nikolaev, Benedikt V. Ehinger, Radha Nila Meghanathan, Cees van Leeuwen

## Abstract

Eye tracking studies suggest that refixations, fixations to locations previously visited, serve to recover information lost or missed during earlier exploration of a visual scene. These studies have largely ignored the role of precursor fixations, previous fixations on locations the eyes return to later. We consider the possibility that preparations to return later are already made during precursor fixations. This would mark precursor fixations as a special category of fixations, i.e., distinct in neural activity from other fixation categories such as refixations and fixations to locations visited only once. To capture the neural signals associated with fixation categories, we analyzed EEG and eye movement recorded simultaneously in a free-viewing contour search task. We developed a methodological pipeline involving regression-based deconvolution modeling, allowing our analyses to account for overlapping EEG responses due to the saccade sequence and other oculomotor covariates. We found that precursor fixations were preceded by the largest saccades among the fixation categories. Independently of the effect of saccade length, EEG amplitude was enhanced in precursor fixations compared to the other fixation categories 200-400 ms after fixation onsets, most noticeably over the occipital areas. We concluded that precursor fixations play a pivotal role in visual perception, marking the continuous occurrence of transitions between exploratory and exploitative modes of eye movement in natural viewing behavior.

## Introduction

In natural viewing, the eyes often return to locations that were visited several seconds before (Yarbus, 1967). Such returns, or refixations, constitute up to 40% of eye movements (Beck, Peterson, & Vomela, 2006; Mannan, Ruddock, & Wooding, 1997; Zelinsky, Loschky, & Dickinson, 2011; Zhang et al., 2022). Their prevalence raises the question of what their role might be, which has made them the subject of intensive eye-tracking research (Gilchrist & Harvey, 2000; Körner & Gilchrist, 2008; Shen, McIntosh, & Ryan, 2014; Tatler, Gilchrist, & Land, 2005; Zelinsky et al., 2011). Results suggest that refixations are aimed at restoration of deficiencies arising in information processing and storage across eye movements. For example, refixations are involved in recovering information that was missed or has become lost during scanning (Gilchrist & Harvey, 2000), updating the representation of a previously visited location (Tatler et al., 2005); rehearsal of a fading memory representation (Meghanathan, Nikolaev, & van Leeuwen, 2019; Zelinsky et al., 2011); or compensating for a premature shift of attention away from the fixation (Gilchrist, North, & Hood, 2001; Peterson, Kramer, Wang, Irwin, & McCarley, 2001).

So far, research has predominantly been centered on the category of refixations while ignoring, arguably, the equally relevant category of *precursor* fixations. These are the first fixations on locations the gaze later returns to, e.g. Fixation 1 in Fig. 1. Precursor fixations until recently have rarely been the focus of eye movement research, and their analysis has yielded inconsistent results. One study showed their duration to be shorter than later refixations. It was concluded that the time at the precursor fixation locations is too short to capture sufficient visual information, and therefore the eyes return (Hooge, Over, van Wezel, & Frens, 2005). Another study, which reanalyzed a large dataset including several experiments with different free-viewing paradigms, found the opposite, namely that precursor fixation locations were fixated longer and were more visually salient than locations the eyes do not return to (Wilming, Harst, Schmidt, & König, 2013). The authors suggested that precursor fixations mark out locations that warrant further scrutiny, while current priority is given to further exploration, causing the eyes to move on and return later for more examination.

**Fig. 1.**
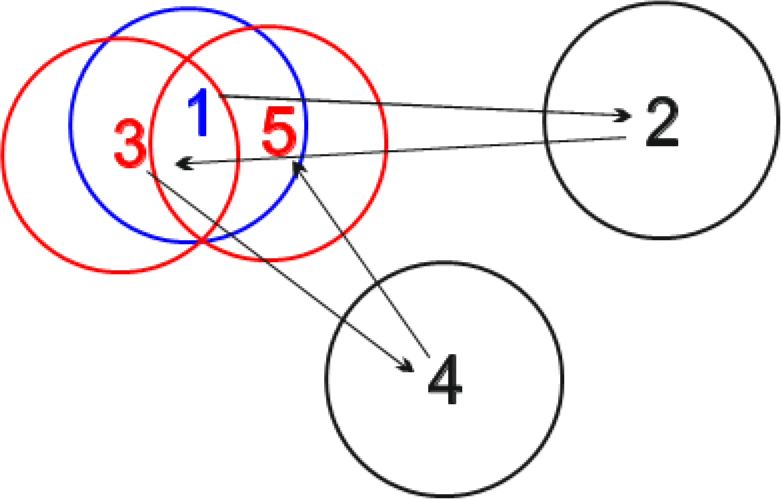
Fixation categories. Numbers indicate fixation rank, i.e., the position in the order of fixations within a trial. Circles represent areas within a radius of 2° of visual angle around the fixation point. We consider fixation 1 as a precursor fixation; fixations 2 and 4 as ordinary fixations; fixations 3 and 5 as refixations. In our previous studies, we did not distinguish between precursor fixations and ordinary fixations. In (Nikolaev et al., 2018), we considered fixations 1, 2 and 4 as ordinary fixations and fixations 3 and 5 as refixations. Similarly, in another previous study (Meghanathan et al. 2020), we considered fixations 1, 2 and 4 as first fixations and fixations 3 and 5 as refixations. Thus, neither of these studies analyzed precursor fixations.

More recently, Zhang and colleagues (2022) analyzed refixation behavior in eight human and animal gaze datasets. Precursor fixations (“to-be-revisited fixations” in the authors’ terminology) were shorter compared to refixations; the difference was particularly pronounced in visual search datasets. The authors suggested that the visual system marks precursor locations for return visits, which utilize additional computational time to improve recognition.

According to Zhang et al. (2022), revisit plans are already being prepared at the time of precursor fixations. Such a bold assertion must be examined against the background of alternative roles that precursor fixations could play. Being initially selected for their saliency, these locations may simply retain this quality during further exploration and hence are likely to continue attracting fixations, regardless of whether the salient information was initially picked up. In this case, there is no distinctive planning, i.e., no return preparation process, at the precursor stage. The eyes return to a location just because it happens to be salient. Or, perhaps the initial gaze did pick up the salient information, but it was forgotten in later viewing so that the gaze had to return to rehearse that information (Zelinsky et al., 2011). If so, again, no specific preparation needs to take place at precursor fixations, insofar as the forgetting is a random process. This may have been the prevailing view of precursor fixations, which may have led them to be ignored. Thus, the role of precursor fixations may have been underestimated and, while little is known about the selection and acquisition of visual information at precursor fixation locations, they may differ from ordinary fixations in ways yet to be understood.

Our current investigation is focused on the neural signals that are associated with precursor fixations. We study precursor fixations using simultaneously recorded eye movement and EEG. Doing so enables unrestricted viewing tasks, which have greater ecological validity than more traditional tasks, in which the eyes are fixed during stimulus presentation. Experimental paradigms with unrestricted eye movement behavior have been used, among others, to study reading (Dimigen, Sommer, Hohlfeld, Jacobs, & Kliegl, 2011), memory encoding and retrieval (Kragel, Schuele, VanHaerents, Rosenow, & Voss, 2021; Nikolaev, Bramão, Johansson, & Johansson, 2023; Nikolaev, Nakatani, Plomp, Jurica, & van Leeuwen, 2011), visual search (Kamienkowski, Varatharajah, Sigman, & Ison, 2018; Körner et al., 2014), perception of natural scenes (Coco, Nuthmann, & Dimigen, 2020; Devillez, Guyader, & Guérin-Dugué, 2015), aesthetic evaluation of art (Fudali-Czyż, Francuz, & Augustynowicz, 2018), and decision making in product valuation (Tyson-Carr et al., 2020). However, none of these studies considered precursor fixations.

By focusing on precursor fixations, the current investigation complements two of our previous studies. Rather than precursor fixations, both studies were dealing with refixations. Like the current one, they used EEG-eye movement co-registration in visual search tasks. In the first (Nikolaev, Meghanathan, & van Leeuwen, 2018), of which the data are also used in the current analysis, participants searched for a collinear contour among a field of randomly oriented Gabor elements. Nikolaev et al. (2018) focused on presaccadic potentials time-locked to the saccade onset (saccade-related potentials, SRPs). Presaccadic potentials indicate saccade planning and preparation, including the shift of attention to the next saccade target (Kovalenko & Busch, 2016; Nikolaev et al., 2011; Wauschkuhn et al., 1998). The presaccadic potential was found to be reduced in amplitude for refixations compared to ordinary fixations over the parieto-occipital regions. The difference may indicate a refixation mechanism operating at the saccade planning stage. The same study also examined postsaccadic potentials. These fixation-related potentials (FRPs), time-locked to fixation onset, reflect initial encoding of visual information at fixation (Kamienkowski et al., 2018; Kazai & Yagi, 2003; Ries, Touryan, Ahrens, & Connolly, 2016). We found a difference in FRP amplitude over the occipital regions about 270 ms after fixation onset between ordinary fixations and refixations. However, this difference was also observed between control fixation pairs (fixations 2 and 4 in Fig. 1), which preserved only the fixation rank (the order of fixations within a trial), but not the actual relationship in the pairs. We interpreted this result as evidence that apart from the effect of fixation order on FRP amplitude, information acquisition at refixations proceeded indistinguishably from ordinary fixations.

In a second study (Meghanathan, van Leeuwen, Giannini, & Nikolaev, 2020), participants searched for multiple target Ts among a field of distractor Ls, all in different orientations. There were 3, 4 or 5 targets and their orientations were to be remembered for subsequent change detection. We compared presaccadic potentials between first fixations and refixations on targets and distractors. For targets only, presaccadic potential amplitude over the occipital regions was higher for first fixations than for refixations. Saccade planning of refixations, therefore, is modulated by top-down factors such as task-relevance.

While both studies identify distinct factors affecting the saccade planning processes immediately prior to a refixation, neither of them provides us with any information regarding precursor fixations. The ordinary fixations in Nikolaev et al. (2018) and first fixations in Meghanathan et al. (2020) are pooled categories, encompassing both precursor fixations and fixations not followed by any refixation (fixations 1, 2, 4 in Fig. 1). The same pooling leaves inconclusive Nikolaev’s et al. (2018) observation that, once the eyes have landed on the revisited location, information acquisition proceeded indistinguishably from ordinary fixations. Thus, an analysis of precursor fixations is crucial in order to understand what happens, not only at the time of the precursor fixation, but also at the time of the refixation.

To examine the neural mechanisms of visual information processing at precursor fixation locations, we compared the amplitude of postsaccadic FRP between three fixation categories: ordinary fixations, refixations, and precursor fixations (Fig. 1). When the return to a precursor fixation location is already prepared at the time of the first visit, we would expect to see a difference between the fixation categories. This difference can, in principle, occur in either of the two known FRP components. The first one is the lambda wave about 100 ms after fixation onset, which is regarded as an early perceptual response of the visual cortex (Dimigen, Valsecchi, Sommer, & Kliegl, 2009; Kazai & Yagi, 1999; Ossandón, Helo, Montefusco-Siegmund, & Maldonado, 2010; Thickbroom, Knezevic, Carroll, & Mastaglia, 1991). The second component is a late parieto-occipital FRP wave after 200 ms, which we have previously attributed to information uptake during refixation behavior (Nikolaev et al., 2018), and which is sensitive to the order of a fixation in a free-viewing trial (fixation rank) (Fischer, Graupner, Velichkovsky, & Pannasch, 2013; Guérin-Dugué et al., 2018; Kamienkowski et al., 2018).

The largest problem of cognitive EEG research in natural viewing behavior is overlapping EEG activity evoked by sequential saccades. Since eye-movement related EEG responses last longer than the respective fixation durations, the EEG response evoked by one saccade does not have time to fade out before the next one and overlaps with it. Moreover, low-level characteristics of eye movement may affect EEG, such as the effect of saccade size on lambda wave amplitude (Dimigen et al., 2009; Kazai & Yagi, 1999; Thickbroom et al., 1991) or the effect of fixation rank on late FRP amplitude (Fischer et al., 2013; Guérin-Dugué et al., 2018; Kamienkowski et al., 2018; Nikolaev et al., 2018). Both types of effects may be systematic and may distort, mask, or be confused with the effects of the cognitive conditions on EEG (Dimigen et al., 2011; Nikolaev, Meghanathan, & van Leeuwen, 2016).

In our two previous refixation studies described above, we solved this problem by restricting comparison between conditions to eye-movement related EEG epochs that were matched on all relevant eye movement characteristics (Dias, Sajda, Dmochowski, & Parra, 2013; Nikolaev et al., 2016). Matching on eye-movement characteristics balances their effects on EEG and thus enables robust comparison between conditions. However, matching involves data loss, because EEG epochs from non-matching eye movement characteristics are excluded from the analysis. Moreover, matching based on absence of statistically significant differences on some eye movement characteristics does not prevent trend contributions from those characteristics (see Dimigen and Ehinger (2021) for more detailed discussion).

Another possible solution to consecutive saccades evoking overlapping EEG activity is based on multiple regression. This technique is similar to the traditional mass-univariate modeling of event-related potentials (ERPs), which was used to adjust for multiple simultaneous discrete and continuous covariates (Smith & Kutas, 2015). However, mass-univariate modeling cannot account for the fact that the degree of overlap in EEG signals varies with the variable fixation durations of successive eye movements in unrestricted viewing behavior. This is because mass-univariate modeling requires segmentation of EEG data relative to a single event (Smith & Kutas, 2015) and has no mechanism to account for EEG activity related to preceding or following events. This was also a shortcoming in our earlier study, where we used generalized additive mixed models (GAMM) of EEG epochs with both experimental conditions and eye movement parameters as covariates (Van Humbeeck, Meghanathan, Wagemans, van Leeuwen, & Nikolaev, 2018).

In multiple regression, the varying temporal overlap can be taken into account by deconvolution modeling (Coco et al., 2020; Cornelissen, Sassenhagen, & Võ, 2019; Dandekar, Ding, Privitera, Carney, & Klein, 2012; Dimigen & Ehinger, 2021; Ehinger & Dimigen, 2019; Guérin-Dugué et al., 2018; Kristensen, Rivet, & Guérin-Dugué, 2017; Litvak, Jha, Flandin, & Friston, 2013). This technique estimates unknown isolated neural responses from the measured EEG and the latencies of experimental and eye movement events, in order to correct for overlapping eye movement effects on EEG. Deconvolution modeling also allows us to correct for the nonlinear effects of various eye movement covariates, such as fixation duration and saccade size and angle. This is achieved by a combination of linear deconvolution with nonlinear spline regression, as used in the generalized additive model (GAM) (Wood, 2017). Deconvolution modeling generates beta coefficients, which represent partial effects of the predictors of interest and correspond to subject-level ERPs in traditional ERP analysis. These beta coefficients are then adjusted for the eye movement covariates included in the model. The adjusted coefficients allow a robust comparison of fixation-related EEG across experimental conditions. Motivated by recent successful examples of deconvolution modeling in face perception (Gert, Ehinger, Timm, Kietzmann, & König, 2022) and episodic memory formation (Nikolaev et al., 2023), in the current study we apply this technique to investigate typical categories of refixation behavior.

## Methods

From the dataset used in this study we previously analyzed EEG in stimulus conditions (Van Humbeeck et al., 2018) and also compared EEG related to refixations and ordinary fixations, but not precursor fixations (Nikolaev et al., 2018).

### Participants

23 healthy adults (two male) took part in the experiment. Data from two participants were removed: one because of problems during eye movement recording and another because of excessive EEG artifacts. The mean age of the remaining 21 participants was 21.6 (range = 18-33) years. All participants gave written informed consent. The study was approved by the Ethics Committee of the Faculty of Psychology and Educational Sciences of KU Leuven and was carried out in accordance with the Declaration of Helsinki.

### Stimuli and procedure

Gabor patches of approximately 0.3-0.4° of visual angle were randomly placed with a mean distance of 0.7° between them in large displays of 30 x 30° at a viewing distance of 55 cm. In half of the trials, at a random location in the display, seven patches formed a contour because their orientation was aligned +/− 25° with the neighboring patches (contour-present trials). In the other half of the trials, the orientation of all patches was random (contour-absent trials).

Participants initiated a trial by pressing the space bar on the computer keyboard. At the beginning of a trial a fixation cross was presented for a random duration between 0.5 and 1 s. Next, a search display was presented for 8 s. Contour-present or contour-absent displays were presented in random order. Participants searched for a contour until the search display disappeared. During a subsequent 5-s response interval, participants indicated the presence or absence of the contour by pressing “p” or “q” keys of the computer keyboard. A feedback screen indicated whether the response was correct. 120 contour-present and 120 contour-absent trials were presented. The trials were organized in 6 blocks of 40 trials with two-minute breaks between blocks. A short practice session preceded the experiment.

### Eye movement recording

The display size necessitates the use of eye movements to search for a contour. Eye movements were recorded with a desktop version of the EyeLink 1000 eye tracking system (SR Research Ltd., Ontario, Canada). The sampling frequency was 250 Hz. A chinrest stabilized participant’s head. A 9-point calibration was performed before each block and whenever it was needed during the block, e.g., if participants occasionally moved their head away from the chinrest. The mean number of calibrations per experiment across participants was 17.7 (range 8-32, SD=7.9). A maximum of 2° error margin between calibration and validation was allowed. The space bar press at the beginning of the trial triggered a drift correction, which allowed tracking errors to be kept within 2°.

### EEG recording

EEG was recorded at a sampling rate of 250 Hz using a 256-channel Geodesic Sensor Net (EGI, a Philips company, Eugene, OR, USA). The net included electrodes for recording the vertical and horizontal electrooculogram. The recording reference was Cz. The EEG was filtered online with an analog high-pass filter of 0.1 Hz and a 100-Hz low-pass filter. TTL pulses were sent through a parallel port from the stimulus presentation computer to the eye tracking and EEG systems. The synchronization of EEG and eye movement recordings was performed offline using the EYE-EEG extension (Dimigen et al., 2011) for EEGLAB (Delorme & Makeig, 2004).

### Selection of refixations, precursor fixations and ordinary fixations

We only analyzed contour-absent trials with correct responses, because only in these trials could visual search reasonably be considered to last the full 8 s duration of the trial. In the contour-present trials, search happened only in initial intervals of uncertain lengths, as the discovery of the contour at any moment terminates search and initiates response preparation. Since a contour, if present, invariably received its first fixation within 3 s (Van Humbeeck et al., 2018), the search was likely too short and the number of fixation-related EEG epochs within it insufficient for reliable EEG analysis.

Fixations and saccades were detected in the gaze data using the velocity-based algorithm for saccade detection (Engbert & Mergenthaler, 2006) of the EYE-EEG extension. In each trial, we identified precursor fixations and refixations within a sequence of fixations (Fig. 1). To identify precursor fixations, we selected all fixations that later receive refixations. A refixation was defined as a fixation within a radius of 2° of visual angle from such a previous fixation. A 2° criterion has repeatedly been used in refixation studies (Anderson, Bischof, Laidlaw, Risko, & Kingstone, 2013; Gilchrist & Harvey, 2000; Solman, Allan Cheyne, & Smilek, 2011). It assures that precursor fixation and refixation locations overlap on the fovea. If a refixation occurred within 2° from two or more close (<2°) precursor fixations, we scored it as a refixation only once. We did not consider as refixations any subsequent fixations prior to leaving the 2° range. If a chain of two or more consecutive refixations occurred, we only kept the first precursor fixation and the immediate refixation, and refrained from assigning these epithets to any item in the chain of refixations. We thus avoided any fixations having both a precursor and refixation label and hence any ambiguity in assigning fixation-related EEG to precursor fixations or refixations. Note however that several refixations can still have the same precursor fixation, so the number of refixations was slightly higher than that of precursor fixations. We also removed fixations immediately preceding a refixation because these contain preparatory EEG activity specific for refixations (Nikolaev et al., 2018).

Finally we selected ordinary fixations, which are fixations at locations to which the eyes never return. Ordinary fixations are ones that are neither refixations nor precursor fixations. They may be considered as fixations “without history”, and are likely to contribute less to oculomotor memory than the other fixation categories. They serve as a reference for categories relating to refixation behavior, and have previously been used for this purpose in our (Nikolaev et al., 2018) and other studies (“non-return fixations” in Zhang et al. 2022).

Fixations not included in any of the selection categories were assigned to the leftover category of ‘other’ fixations. As explained in the “Deconvolution” section below, this category is technically needed for deconvolution to correct for *all* overlapping effects of eye movements on the EEG. It contains fixations outside of the areas of interest, but also fixations that could not be unambiguously designated as ordinary fixations, precursor fixations, or refixations. This included, for instance, chains of fixations on the same location, which may equally qualify as refixation of the previous or as precursor for the next fixation. While ‘other’ fixations form a heterogeneous bunch needed only for correct operation of the deconvolution modeling, the ordinary fixations have a definite functional significance as an intrinsic part of refixation behavior.

Using the EYE-EEG extension, for the current eye movement, we extracted fixation duration, X and Y fixation position (the mean horizontal and vertical screen coordinates of a fixation, respectively, in pixels), size and angle of the saccade incoming to the current fixation. Since differences in viewing behavior may occur not only at the current fixation but also at the next one, we also extracted the size and angle for the outgoing saccade, as well as the duration of the immediately following fixation (only if the following fixation was of the ‘ordinary’ or ‘other’ category).

### EEG cleaning

Data processing consisted of two main parts: cleaning the EEG data, and modeling the overlapping effects of sequential eye movements and their covariates on the EEG. Cleaning was performed on continuous EEG, which is an essential requirement for the application of deconvolution modeling with time expansion for correction of overlapping effects, as these may be produced by all the eye movements and other events in the experiment.

We analyzed 148 of the 256 electrodes (Fig. 3D): 108 electrodes close to the cheeks and neck were removed because they often had poor contact due to the long hair of our mostly female participants, and showed strong muscle artifacts. For cleaning we used functions from the EEGLAB toolbox for MATLAB. First, we applied the *pop_cleanline* function, which removes power line noise from EEG using multi-tapering, and a Thompson *F*-statistic. This method uses a 4-s sliding window (with 1-s step), in which a frequency-domain regression estimates the amplitude and phase of the sinusoidal signal at the power-line frequency (50 Hz) and its harmonics. The fitted signal is subtracted from the EEG. Then we applied *clean_artifacts*, which removes flat-line channels, low-frequency drifts, noisy channels, and short-time bursts. To remove transient or large-amplitude artifacts this function uses artifact subspace reconstruction (ASR). It is an automatic, subspace-based method, which compares the structure of the artifactual EEG activity to that of known artifact-free reference data (Mullen et al., 2015). Artifact-free data from a given EEG recording are obtained by automatically selecting low-variance EEG sections. ASR then calculates a clean subspace based on this activity. Next, ASR projects 1-s sliding windows into this clean subspace, which allows for reconstruction of an artifact-reduced EEG signal. Instead of using such an interpolated signal directly, ASR takes deviations of the reconstructed signal to the raw data as an indication of noise-contamination and marks these portions for rejection. The efficiency of ASR for artifact removal crucially depends on the ASR parameter that defines the tradeoff between removing non-brain signals and retaining brain activities. We set the ASR parameter to 20, which was found to be optimal in a dedicated study (Chang, Hsu, Pion-Tonachini, & Jung, 2020). On average, this procedure removed 10.5 (SD=7.3) channels per participant, but entire EEG segments were not removed, given the crucial requirement to use continuous EEG for deconvolution modeling.

We removed ocular artifacts using the OPTICAT function (Dimigen, 2020). This function is designed to remove all types of ocular artifacts in unrestricted viewing behavior (e.g., eyeball rotation, blinking), with a particular focus on saccadic spike activity. Like most artifact removal methods, this function is based on independent component analysis (ICA), but requires preparation of the input EEG, which isolates influences of oculomotor artifacts better than ICA training on unprepared EEG data. The preparation involves high-pass EEG filtering at 2 Hz but no low-pass filtering, to preserve the high-frequency (> 40 Hz) components of myogenic activity, which are characteristic of the saccadic spike potential (Keren et al. 2010). A key step of the OPTICAT function is overweighting the contribution of saccadic spike activity in the EEG input to ICA. This improves artifact correction, particularly of saccadic spike activity. Overweighting of the filtered data was achieved by cutting 30-ms EEG segments (−20 to +10 ms) around saccade onsets as identified by eye tracking, and repeatedly appending these segments to the EEG. Then, ICA was trained on these overweighted data, and the obtained ICA weights were transferred to the unprepared data. The variance of each independent component during saccade and fixation intervals was calculated. Components with greater variance during saccade than fixation intervals are more likely to reflect ocular artifacts. The ratio between the mean variance during saccade and fixation intervals was computed. If the ratio exceeded 1, the component may already reflect ocular artifacts, but in effect, the ratio threshold was set slightly higher, at 1.1, as this is considered optimal for reducing misclassifications (Plöchl, Ossandón, & König, 2012). The ratio of each independent components was checked against this threshold. If the ratio exceeded this threshold, the corresponding components were considered to be saccade-related and were removed. On average, 4.2 (range 2-7, SD=1.37) saccade-related components per participant were removed. (Fig. S1 shows maps of saccade-related components intended for rejection and the distribution of variance ratio of all components for a representative participant).

Next, the ICs related to the remaining artifacts were removed with the automatic classifier (*pop_iclabel*) (Pion-Tonachini, Kreutz-Delgado, & Makeig, 2019). The probability ranges of independent components to be removed as artifacts were set as follows: ‘brain’ 0-0, ‘muscle’ .4-1, ‘eye’ .9-1, ‘heart’ .05-1, ‘line noise’ .4-1, ‘channel noise’ .4-1, ‘other’ .4-1. On average, 84.7 (range 50-107, SD = 15.6) components were removed, leaving about 63 ‘brain’ and mixed components per participant. Most of the removed components were classified as ‘channel noise’ (27.3) or ‘other’ (51.7) (Supplementary Table S1). The EEG was then recreated without these components. Finally, EEG was filtered with a low cut-off of 0.1 Hz (−6 dB at 0.05 Hz) and with a high cut-off of 30 Hz (−6 dB at 30.05 Hz) using the *pop_eegfiltnew* function with default settings. The high cut-off of 30 Hz was sufficient for the FRP components of interest, because lambda activity is most prominent in the upper-theta and alpha bands (6–14 Hz) (Dimigen et al., 2009; Nikolaev et al., 2016; Ossandón et al., 2010), and the frequency of the late FRP component was even lower, as seen in Fig. 3A. EEG was re-referenced to average reference. The channels previously removed by *clean_artifacts* were interpolated with spherical spline interpolation.

### Deconvolution

Due to the non-uniform distribution of fixation durations, effects of sequential eye movements in natural viewing behavior on EEG are systematic and may confound effects of experimental conditions. Moreover, low-level oculomotor characteristics, such as fixation duration, X and Y positions of a fixation on the screen, the size and angle of saccades, *per se* may affect fixation-related EEG (Cornelissen et al., 2019; Dimigen et al., 2011; Nikolaev et al., 2016). In addition, the EEG response evoked by the onset of the stimulus screen may distort the following fixation-related potentials (Dimigen et al., 2011; Gert et al., 2022).

To eliminate these effects, we used the deconvolution approach implemented in the Unfold toolbox for MATLAB (Ehinger & Dimigen, 2019). This approach involves regression-based estimation of ERP (for an in-depth review see Smith & Kutas, 2015). The toolbox performs a regression-based EEG analysis that includes mass univariate modeling, linear deconvolution modeling, and generalized additive modeling. As a result, it computes the partial effects (i.e., the beta coefficients or “regression ERPs, rERPs”) of predictors of interest, adjusted for all other covariates. The analysis with the Unfold toolbox consists of four major steps, which are described in detail in (Ehinger & Dimigen, 2019) and are illustrated with respect to different data types in (Dimigen & Ehinger, 2021). As the first step, we specified the regression model and generated the design matrix. According to the Wilkinson notation, the model formula was defined as follows:

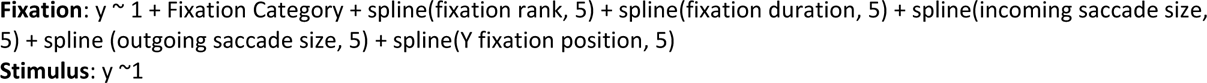

The formula indicates that we considered multiple effects of the fixation event and the stimulus onset event. Specifically, for fixations, we considered as covariates the fixation onset (y ∼ 1, i.e., the intercept) and Fixation Category as a categorical predictor with four levels: precursor fixations, refixations, ordinary fixations, and ‘other’ fixations. We used treatment (dummy) coding with ‘other’ fixations as the reference level (intercept). As a result, the first three levels (precursor fixations, refixations, ordinary fixations) were coded as differential FRPs relative to ‘other’. (Note that in a linear model, it does not matter which level of the categorical predictor is used as the reference level, as marginal predictions are interpreted). The ‘other’ fixations category is technically necessary to ensure high quality correction in any deconvolution model involving successive fixations (e.g., Gert et al., 2022; Nikolaev et al., 2023). The ‘other’ fixations allowed us to account for the overlapping effects on continuous EEG produced at the latencies of ‘other’ eye movements. The responses to ‘other’ eye movements could spill over to the responses elicited by the eye movements of interest, because they often interleaved between precursor fixations, refixations, and ordinary fixations. To control for overlapping effects of successive eye movements, we needed to consider all fixations that occurred in the experiment, without exception. Thus, even though ‘other’ fixations are not specific for refixation behavior, they constitute an essential level of the predictor variable ‘Fixation Category’.

Eye movement covariates were selected for inclusion in the model based on differences in respective eye movement characteristics between fixation categories, as described below in the section ‘Selection of eye movement covariates for deconvolution correction’.

Since we assumed that the selected eye movement covariates have nonlinear effects on EEG, we modeled them with a basis set of five spline predictors. Automatic procedures for estimating the number of splines are currently not available. We followed the recommendation of Dimigen & Ehinger (2021) and set the number of splines to five, thereby limiting the “wiggliness” or un-smoothness of the nonlinear relationship. Spline knots were placed on the participant-specific percentiles of the covariates (i.e., on the 3 (N_splines_-2 = 5-2) quantiles). We did not include interaction between splines in this dataset, as these would have required fitting 2D-splines, raising the number of predictors beyond feasible for the current dataset. We modeled the potentials evoked by the stimulus events using only a single waveform for all images (y ∼ 1, i.e., the intercept).

In a second step, the design matrix was time-expanded in the time window between −200 and +500 ms around fixation and stimulus onset events, using the function *uf_timeexpandDesignmat* of the Unfold toolbox, in which 175 ‘local’ time points around each event are added as predictors (175 points due to a sampling rate of 250 Hz and 700 ms duration). The total number of predictors before time-expansion was 25. This number is the sum of one intercept, four categorical predictors relating to the fixation categories, and five eye movement (and fixation rank) predictors modeled with five splines (i.e., 5*N_splines_-1 = 20, per spline we remove one parameter, similar to when modeling the categorical predictor in the presence of an intercept). Per event type, the time expansion takes the respective mass-univariate design-matrix (number of events x 25 predictors) and generates a new design matrix applicable to the continuous EEG (see Fig. 4 in Dimigen and Ehinger (2021) for an illustration of time-expansion). Each event is modeled by a set of time-lagged impulse response functions, one for each time lag, – weighted by its respective mass-univariate design-matrix value. These values are pasted into the new design matrix around the onset of the respective event.

Because the new design matrix is defined for all time points of the continuous EEG, we estimate only a single linear model instead of estimating one linear model per time-point, as in the mass univariate approach. This allows us to simultaneously estimate all stimulus and fixation betas, accounting for their potential overlap effects. The time-expanded design matrix spanned the duration of the entire EEG recording. It had 4375 (25 predictors x 250 Hz x 0.7 s) columns. The number of rows varied across participants because of the different number of events (on average 1,343,503 (range 1,076,301-1,773,579, SD=206,900) rows per participant). Thus, for 4375 parameters, we obtained approx. 1,340,000 data points. Since the number of data points per parameter exceeds the number of parameters by two orders of magnitude, the model is not over-parameterized.

In the third step, we excluded irrelevant intervals from modeling by setting entire rows of the time-expanded design matrix to zero, using the function *uf_continuousArtifactExclude*. Filling with zeros preserved the timing of continuous EEG. Particularly, we removed from the model the inter-trial intervals, the breaks between blocks, the contour-present trials, the trials with incorrect responses, the trials without refixations, and bad eye-tracking intervals. Then, the deconvolution model was fitted (*uf_glmfit*) for each of the 148 electrodes using the iterative Least Squares Minimal Residual algorithm (Fong & Saunders, 2011) for sparse design matrices.

In the fourth and final step, we reconstructed averaged EEG waveforms (“regression-based fixation-related potentials”, rFRPs) from the beta coefficients of the model for the three fixation categories for each participant using the functions *uf_predictContinuous* and *uf_addmarginal*. In principle, the raw beta coefficients could be interpreted and compared statistically, since they represent the “pure” effects of the predictors. These effects may be considered as a rough analog of difference waves relative to a reference level. Accordingly, the waveforms of the “pure” effects have highly wiggly shapes, similar to ERP difference waves. This complicates the comparison of deconvolutional and traditional ERP results. We addressed this issue by adding a model intercept to the difference wave. But since we used nonlinear modeling with splines, there was no clear reference level for the nonlinear basis set (the set that transforms predictor values by weighting beta coefficients). Therefore, we estimated all nonlinear effects at the corresponding mean predictor levels and added for each fixation category these estimates to the beta coefficients. This ensured that all fixation categories are presented at the same estimated predictor levels. The resulting waveforms were comparable to the participant-level averages in traditional ERP analyses.

### Statistical analysis

Deconvolution was performed for each participant separately. In principle, we could compute group-statistics across individual partial-R². However, at present, partial-R² can only be evaluated for a whole epoch, not on individual time-points. To perform the statistical analysis at the group level, we used a repeated-measures ANOVA on the mean FRP amplitudes in the time windows described below. The Huynh-Feldt correction for p-values associated with two or more degrees of freedom was applied in case of sphericity violation. The Newman-Keuls test was used for post-hoc analyses.

To perform the statistical analysis on the group level we estimated the FRP amplitude in unbiased time windows based on the grand average and *a priori* regions of interest (ROIs). Since the number of *a priori* tests is typically small, the issue of multiple comparisons for them is minor, which increases the statistical power of the tests. Moreover, they are more likely to detect narrowly distributed effects that occur across a small number of time points and electrodes (Groppe, Urbach, & Kutas, 2011). We selected eight ROIs over the left and right hemisphere, namely, frontal, central, parietal and occipital brain regions, which were defined around landmark electrodes of the International 10-20 System of Electrode Placement: F3, F4, C3, C4, P3, P4, O1, O2 (the inset in Fig. 3D).

For each ROI, we averaged amplitudes over one central and six surrounding electrodes. This approach involved 56 (=7 x 8) electrodes, symmetrically and systematically distributed over the head. Such an approach is generally suggested for high-density electrode montages (Dien & Santuzzi, 2005). Based on visual inspection of the grand-averaged FRPs we selected two time windows. The first, lambda time window consisted of the 30-130 ms interval after fixation onset, which was chosen as the 100 ms window centered on the lambda peak latency of 80 ms, a latency which was found by averaging across all conditions and participants. The second, late, time window was chosen to include the most pronounced FRP negativity over the occipital regions 200-400 ms after fixation onset.

For the FRP amplitude in the lambda time window, we considered only the occipital ROIs (OL and OR) because here the lambda activity is maximal (Kazai & Yagi, 2003; Thickbroom et al., 1991; Yagi, 1979). Therefore, the ANOVA had two factors: Fixation category (Precursor Fixation, Refixation, Ordinary Fixation) and Hemisphere (left, right). For the late time window, we considered all 8 ROIs, and therefore the ANOVA had three factors: Fixation category (Precursor Fixation, Refixation, Ordinary Fixation), ROI (frontal, central, parietal, occipital), and Hemisphere (left, right). The statistical analyses were performed with STATISTICA 10 (StatSoft Inc., Tulsa, OK, USA) and R version 4.2.0 (2022).

## Results

### Eye movement results

There were on average 205 (range 130-270, SD=39.9) precursor fixations, 219 (range 138-286, SD=45) refixations, 1680 (range 1266-2347, SD=284.5) ordinary fixations, and 402 (range 246-597, SD=103) ‘other’ fixations per participant.

For fixation duration, a repeated-measures ANOVA with the factors of Fixation category (precursor fixations, refixations, ordinary fixations) and Fixation order (current vs following fixation) showed an effect of Fixation category (F(2, 40) = 6.8, p = .005, ε = .87), Fixation order (F(1, 20) = 10.9, p = .004), and an interaction between them (F(2, 40) = 19.5, p < .001, ε = .92) (Fig. 2A). Post-hoc tests revealed no difference in duration between fixation categories of the current fixation (all p > .4), but differences in duration between all categories of the following fixation (all p < .002): average duration of fixations following refixations was longest while those following precursor fixations were shortest (Fig. 2A). Moreover, precursor and ordinary fixations had longer durations than fixations following them (all p < .04), while the duration did not differ between refixations and their following fixations (p = .22). For saccade size, an ANOVA with the same design showed an effect of Fixation category (F(2, 40) = 14.9, p < .001, ε = .98), Fixation order (F(1, 20) = 11.3, p = .003), and an interaction between them (F(2, 40) = 47.4, p < .001, ε = .86) (Fig. 1B). Post-hoc tests revealed that incoming saccades were largest in size for precursor fixations and smallest for refixations (all p < .001). Outgoing saccades were largest for ordinary fixations and smallest for precursor fixations (all p < .02). Among fixation categories, for precursor fixations had the largest incoming saccade size and the smallest outgoing saccade. Moreover, saccades incoming to precursor fixations were larger than outgoing saccades (p < .001). In contrast, saccades incoming to refixations and ordinary fixations were smaller than outgoing saccades (all p > .04). For saccade angles, a circular ANOVA (the R package ‘Circular’, v0.4-93; (Lund & Agostinelli, 2017)) on data pooled across participants showed no significant differences between fixation categories for incoming (F(2, 44181) = 1.1, *p* = .3) or outgoing saccades (F(2, 44181) = 0.5, p = .6) (Fig. 2G). Fixation positions were compared between fixation categories separately for X and Y positions. No effect of fixation categories on the X fixation position was found (F(2, 40) = 0.7, p = .47). For the Y position, we found an uncorrected effect of fixation categories (p = .025) which, however, lost significance after Huynh-Feldt correction: (F(2, 40) = 4.04, p = .052, ε = .56) (Fig. 2E). Nevertheless, we performed a post-hoc test and found that the Y position of ordinary fixations differed from that of refixations (p = .04) and precursor fixations (p = .02) (as expected, positions of refixations and precursor fixations were about the same (p = .83)).

**Fig. 2.**
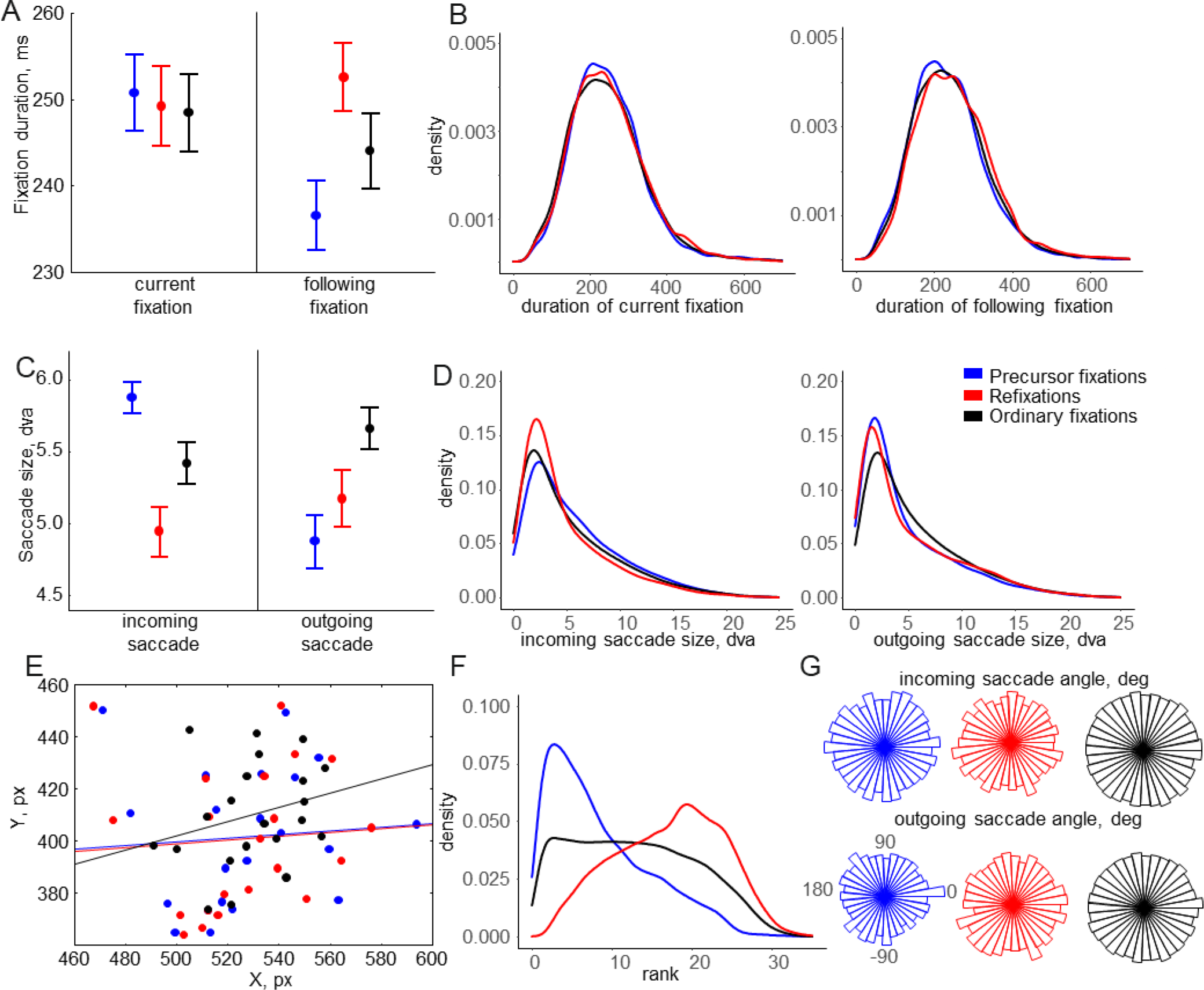
Eye movement results for three fixation categories. A: Duration of the current and following fixations (mean-error plots, with error bars indicating the standard errors of the means across 21 participants). B: Probability density estimation of the corresponding fixation durations. C: Size of the incoming and outgoing saccades. D: Probability density estimation of the corresponding saccade sizes. E: Distribution of mean fixation positions on the screen for each participant (each dot represents one participant). The lines indicate the linear fits of the distributions, emphasizing the difference in the Y position of ordinary fixations. F: Probability density estimation of the fixation rank within an 8-s trial. G: Distribution of angles of the incoming and outgoing saccades.

The probability density plot of fixation rank shows that precursor fixations and refixations were distributed unequally in time, skewing toward the left and right edges of a trial, respectively (Fig. 2F). Precursor fixations and refixations were separated by, on average, 9 (SD = 1.2, range 7-10.9) intervening fixations.

Our results show changes in oculomotor behavior characteristic of precursor fixations: a decrease in outgoing saccade size and duration of the following fixation, which fails to occur for refixations and ordinary fixations. Moreover, size of incoming and outgoing saccades and Y fixation positions differ between fixation categories. These differences may confound FRP effects associated with fixation categories and should be corrected before further analysis.

### EEG results

#### Selection of eye movement covariates for deconvolution correction

Differences between eye movements during unrestricted viewing could have an effect on EEG confounding effects of the conditions of interest (Dimigen et al., 2011; Nikolaev et al., 2016). To correct these confounds, we applied deconvolution modeling. The choice of eye movement covariates and other predictors to be corrected by deconvolution is not a simple matter. The desire to include as many of these predictors as possible is faced with theoretical (e.g., collider bias) and practical (e.g., overfit) limitations. The former cannot easily be resolved without explicit causal modeling (Rohrer, 2018). But overfitting can be diagnosed in extreme cases by investigating the number of iterations of the solver or, more generally, by cross validation, and can be remedied by simplifying the model, for instance, by reducing the number of predictors. Therefore, we chose a data-driven strategy for selecting predictors using a statistical test for effect in the conditions of interest and using only those for which the effect was significant.

Specifically, our inclusion criterion for deconvolution correction of eye movement covariates was the presence of a difference between fixation categories, as reported in the section “Eye movement results”. We included the fixation rank, i.e., the position in the order of fixations within a trial, because it differed between precursor fixations and refixations by definition (Fig. 2F). Moreover, it is known that the amplitude of fixation-related potentials during a trial with multiple eye movements depends on the fixation rank (Fischer et al., 2013; Guérin-Dugué et al., 2018; Kamienkowski et al., 2018; Nikolaev et al., 2018). We included the size of incoming and outgoing saccades, which differed between fixation categories (Fig. 2C), but we did not include saccade angles, since they did not differ. We also considered the X and Y screen coordinates of the fixation locations, as it has previously been reported that the fixation location influences FRP (Cornelissen et al., 2019). We found a trend for the difference between fixation categories for Y but not for X fixation position (Fig. 2E), and therefore only included the Y position. Finally, we included the duration of the current fixation. Though this duration did not differ between fixation categories, it is indicative of the onset latency of outgoing saccades. The average (across all categories) duration in our task was 247 ms, and thus most of subsequent eye movements occurred in the middle of the late time window (200-400 ms after fixation onset). Saccadic-related potentials evoked by following eye movements may interfere with ongoing FRPs in this time window, especially given that outgoing saccades differ in size between conditions.

It should be noted that we currently only tested the eye movement characteristics for effects on the means. This strategy comes with limitations. Due to a possible non-linearity in the EEG response to eye movement covariates, differences may be present also in the distribution, which should also be tested. In a future scenario, the selection could be based on a priori considerations based on the availability of a theory or preliminary studies.

#### Selection of a baseline interval

In EEG research, an ideal baseline interval should contain no traces of evoked or induced EEG activity. Therefore, in a stimulus-response paradigm with fixed eye movements, the preferred baseline interval is usually the one before the stimulus onset. The EEG co-registered with unrestricted eye movements, however, consists of overlapping brain responses evoked by consecutive saccades. This makes it difficult to find a baseline interval that would be neutral in relation to the fixation-related EEG epoch. We have previously described several options for locations of the baseline interval for fixation-related epochs during unrestricted viewing (Nikolaev et al., 2016). For example, a common baseline could be selected before the onset (stimulus presentation) of a trial involving multiple eye movements, for application to each fixation-related epoch of that trial; alternatively, an individual baseline could be selected from some interval around a saccade and applied to the rest of the fixation-related epoch. All options have their pros and cons, and the final choice of baseline depends on the goals and design of the study. However, it is obvious that the baseline interval used should not be affected by the experimental conditions of interest, otherwise baseline correction would transfer to the baseline FRP difference between conditions for the rest of the fixation-related EEG. Consequently, in this study we checked for a difference between the fixation categories in the two candidate baseline intervals.

The first candidate baseline interval was chosen from −200 to −100 ms before fixation onset; an interval that precedes and thus excludes effects of saccade execution. The second candidate baseline interval was chosen from 0 to 20 ms after fixation onset. This interval was considered already free from the influence of saccade preparation and execution, while the perception-related EEG response has not yet begun (Rama & Baccino, 2010). We applied a repeated-measures ANOVA with factors of Fixation category (Precursor fixation, Refixation, Ordinary fixation), ROI (frontal, central, parietal, occipital), and Hemisphere (left, right) on the mean amplitude in these intervals of the deconvolved but not baseline-corrected FRPs. For the first interval, we found an interaction of Fixation category with ROI (p = .03), which, however, lost significance after Huynh-Feldt correction: (F(6, 120) = 2.4, p = .1, ε = .38), and a triple interaction of Fixation category x ROI x Hemisphere (p = .02), which also lost significance after Huynh-Feldt correction: (F(6, 120) = 2.7, p = .06, ε = .43). For the second interval, we found no main effect or interactions involving Fixation category. Although the interactions for the first interval were not significant, they may suggest a trend for the effect of Fixation category on EEG. Therefore, in all our further EEG analyses we opted for the second interval as our baseline interval. It should be noted that choosing a baseline interval for which no differences are found between experimental conditions is tantamount to accepting a null-hypothesis and is therefore only a heuristic. More advanced procedures, such as baseline regression (Alday, 2019), are not currently compatible with deconvolution overlap correction.

#### The effect of fixation categories on FRP

Fig. 3A shows the beta coefficients (FRPs) obtained after deconvolution and adding covariates to the fully adjusted model, grand-averaged across 21 participants in the three fixation categories. The shape of the waveforms is similar to that expected in traditional FRPs time-locked to the fixation onset and is characterized by a prominent lambda wave over the occipital regions. To examine the effect of fixation categories on these FRPs, we applied an ANOVA on the mean amplitude in the lambda and late time windows.

**Fig. 3.**
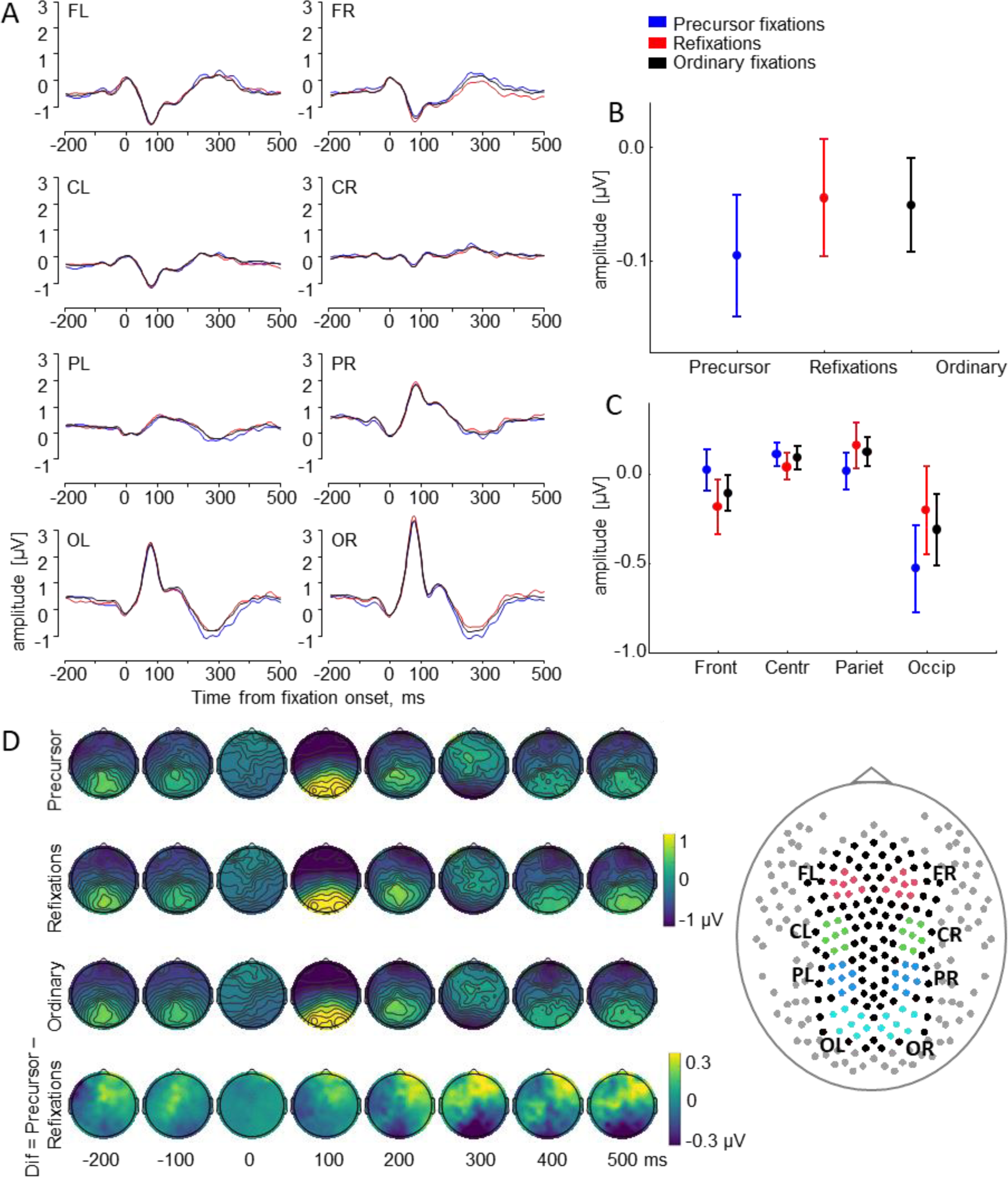
Fixation-related potentials (regression betas) time-locked to the fixation onset for three fixation categories. The potentials are grand-averaged (N=21) and baseline-corrected at 0-20 ms after fixation onset. A: Fixation-related potentials for 8 ROIs. B: Mean amplitude in the interval 200-400 ms after fixation onset, indicating a main effect of fixation category. C: Mean amplitude in the interval 200-400 ms after fixation onset, indicating an interaction between fixation category and ROI. Error bars indicate standard errors of the means across 21 participants. D: Momentary maps over 148 electrodes for the three fixation categories, as well as a difference map obtained by subtracting FRP of refixations from those of precursor fixations. The inset on the right indicates the locations of all 256 recorded electrodes. Removed electrodes are shown in grey and remaining ones either in black or in different colors according to regions of interest (ROIs): left and right frontal (FL, FR), central (CL, CR), parietal (PL, PR), occipital (OL, OR).

In the lambda time window (30-130 ms after fixation onset), there was no main effect of Fixation category on the FRP amplitude (p = .49), but there was a main effect of Hemisphere with significantly higher FRP amplitude over the right than over the left occipital ROIs (F(1, 20) = 12.5, p = .002). There was no interaction between Fixation category and Hemisphere (p = .9).

In the late time window (200-400 ms after fixation onset), the FRP amplitude for precursor fixations was visually different from other fixation conditions, with a maximal difference at approximately 250 ms after the fixation onset (Fig. 3A). This was supported by a significant main effect of Fixation category (F(2, 40) = 3.3, p = .048, ε = 1) (Fig. 3B). Post-hoc tests showed a difference in FRP amplitude between precursor and ordinary fixations (p = .045) and a near-significant difference between precursor fixations and refixations (p = .06). The difference for precursor fixations was most prominent over the occipital ROIs, as evidenced by the interaction between Fixation category and ROI (F(6, 120) = 3.7, p = .03, ε = .37) (Fig. 3C). Post-hoc tests revealed a lower FRP amplitude over the occipital ROIs for the precursor fixations compared with refixations (p < .001) and ordinary fixations (p = .015), and no difference between refixations and ordinary fixations (p = .21).

Topographical maps for 148 electrodes showed a positive peak, equally large for all three fixation categories. The peak was narrowly localized over the occipital areas 100 ms after the fixation onset – the lambda wave (Fig. 3D). At 300 ms after fixation onset, a prominent negativity was spread over the parieto-occipital areas and was mirrored over the frontal areas. The area of negativity over the parieto-occipital areas was larger for precursor fixations than for refixations and ordinary fixations. The difference maps in Fig. 3D show the topography of the FRP difference between precursor fixations and refixations, which was largest among the fixation categories. Thus, the FRPs for precursor fixations differed from FRPs for refixations and ordinary fixations over the occipital areas.

The FRP amplitude in the late time window did not differ between refixations and ordinary fixations. To understand better the relationships between these fixation categories, we considered how refixations and ordinary fixations were related to the category of ‘other’ fixations, which were used as the reference level (intercept) in the deconvolution modeling. To this end, we conducted two additional analyses. In the first, we statistically estimated the role of ‘other’ fixations among the fixation categories. In the second, we started from the assumption that the neurophysiological manifestation of the ordinary fixations might be similar to ‘other’ fixations, due to the mixed nature of the brain processes underlying both categories. Thus, we combined ordinary and ‘other’ fixations into one category and ran the deconvolution modeling with the three fixation categories: precursor fixations, refixations, ‘ordinary+other’. The results of these analyses are shown in the Supplementary Fig. S2.

In the first analysis, we extracted the mean FRP amplitude for ‘other’ fixations (intercept) in both time windows and ran the same ANOVAs as in the main analysis, but now with four fixation categories (Fig. S2A). In the lambda time window, we found no main effect and no interaction involving fixation categories (all p > .5). In the late time window, we found an interaction between Fixation category and ROI (F(9, 180) = 2.5, p = .01), which, however, was reduced to marginally significant after accounting for sphericity violation (p = .057, ε = .4). Post-hoc tests revealed a lower FRP amplitude over the occipital ROIs for the precursor fixations compared to all other categories (all p < .02) and no difference between any of the other categories (all p > .46) (Fig. S2B).

In the second analysis, with combined ordinary and ‘other’ fixations, in the lambda time window we found no main effect and no interaction involving fixation categories (all p > .6), as before. In the late time window, we found a significant main effect of Fixation category (F(2, 40) = 4.0, p = .03, ε = 1) (Fig. 3B). Post-hoc test showed a lower FRP amplitude for precursor fixations than for ‘ordinary+other’ fixations (p = .02) and refixations (p = .047). We also found an interaction between Fixation category and ROI (F(6, 120) = 3.4, p = .036, ε = .37). As before, post-hoc test revealed a lower FRP amplitude over the occipital ROIs for the precursor fixations as compared to all other categories (all p < .006) and no difference between the refixation and ‘ordinary+other’ categories (p = .7) (Fig. S2D).

Thus, the additional analyses highlighted the specificity of precursor fixations, which are lowest among the four fixation categories in FRP amplitude over the occipital areas in the late time window. The FRP for ‘other’ fixations did not differ from the FRPs for refixations and ordinary fixations.

## Discussion

We considered the possibility that during precursor fixations, which are fixations on locations to which the eyes return during later visual exploration, the plan to return later is already being prepared. This should mark the category of precursor fixations as different in their neural signal from the categories of refixations and ordinary fixations, which are neither precursor fixations nor refixations. We analyzed co-registered eye movements and EEG from a free-viewing contour-search task, comparing oculomotor and neural properties of precursor fixations with those of refixations and ordinary fixations. Regression-based deconvolution modeling was applied to correct fixation-related potentials (FRPs) for effects of multiple eye movement covariates and for the overlapping effects of successive eye movements naturally occurring in unrestricted viewing.

No difference in effect between fixation categories was obtained for lambda activity, which is known to reflect early perceptual processes at each fixation (Dimigen et al., 2009; Kazai & Yagi, 1999; Ossandón et al., 2010; Thickbroom et al., 1991). Specifically, lambda activity is considered a response of the visual cortex to retinal image displacement following eye movement (Dimigen et al., 2009; Kazai & Yagi, 1999; Thickbroom et al., 1991), which has a sensitivity profile and cortical sources similar to the P1 component in event-related potentials (Kazai & Yagi, 2003).

Differences among fixation categories were obtained, however, for the late FRP. The FRP of precursor fixations differed in amplitude from ordinary fixations and refixations 200-400 ms after fixation onset, most noticeably over the occipital areas. Since this difference was observed after correcting the EEG for eye movement covariates, it cannot be understood as an oculomotor effect. Thus, the effect is likely to reflect distinct sampling of visual information at precursor fixations. This effect would be in accordance with our proposal that already during precursor fixations a return plan is being prepared.

The evoked component affected by fixation categories occurs in our analysis in the late time window. There is evidence that the FRP amplitude in this time window depends on fixation rank. For example, a more negative occipital FRP amplitude for precursor fixations than for refixations in this time window corresponds to the observation by Fischer and colleagues (2013) of a more negative parieto-occipital FRP amplitude for fixations earlier in a trial than for later ones (cf. our Fig. 3A and their Fig. 2A, although they did not analyze this effect statistically). The dependence of FRPs on fixation rank in this time window has also been found in the perception of emotional faces (Guérin-Dugué et al., 2018) and in a free-viewing visual search task (Kamienkowski et al., 2018). This dependence may reflect integrative cognitive processes that become more prominent as search progresses. In the initial analysis of the current dataset (Nikolaev et al., 2018) we already observed an effect of fixation rank in the late time window. However, this effect was not specific to refixations, as it was reproduced in a control analysis with ordinary fixations whose fixation rank matched to that of refixations. In the current analysis, the oculomotor effect of fixation rank on FRP was eliminated by deconvolution, and consequently there was no spurious FRP difference between ordinary fixations and refixations. Thus, the observed FRP difference of precursor fixations from ordinary fixations and refixations represents a true effect of sequential visual sampling on brain activity. It may be related to higher-order processes accompanying visual sampling in the sequence of eye movements. In particular, perception in unrestricted viewing involves sampling, integrating, and accumulating visual information from multiple locations in a scene (Hollingworth & Henderson, 2002; Melcher & Kowler, 2001; Tatler, Gilchrist, & Rusted, 2003). Representations of these locations are stored in visual short-term memory for further use (Higgins & Rayner, 2015; Irwin & Zelinsky, 2002; Prime, Vesia, & Crawford, 2011). The representation and accuracy of short-term memory information are supported by the early visual areas in the occipital lobe (Emrich, Riggall, Larocque, & Postle, 2013; Sligte, Scholte, & Lamme, 2009), consistent with the observed predominance of the fixation category effect over the occipital areas. Thus, the brain activity that distinguishes precursor fixations from refixations and ordinary fixations may be related to memorizing locations in order to return to them later.

The relationship between precursor fixations and refixations could be understood within the concepts of exploitation and exploration. The exploratory mode of eye movement involves broad scanning of a visual scene as the focus of attention moves across new regions. In contrast, the exploitative mode involves lingering on a region and scrutinizing its details, as well as revisiting previously seen regions. Accordingly, exploration is characterized by a wider spatial distribution of eye movements, whereas exploitation is characterized by longer fixation durations (Ehinger, Kaufhold, & König, 2018; Ramos Gameiro, Kaspar, König, Nordholt, & König, 2017; Wilming et al., 2013). Regulation of the balance between these competing brain processes is a fundamental property of adaptive behavior (Cohen, McClure, & Yu, 2007). Its dynamic is manifested in the transitions between exploratory and exploitative viewing modes during scene inspection (Ramos Gameiro et al., 2017) and is associated with the dynamics of holistic and analytic perception of complex visual stimuli (Fitousi, 2020; Navon, 1977; Van Leeuwen & Bakker, 1995). The sudden appearance of a visual stimulus initiates exploratory behavior with large saccades and short fixations at the beginning of a free-viewing trial. Later in the trial, saccade size decreases and fixation duration increases, reflecting the transition from an exploratory (ambient) to an exploitative (focal) viewing mode (Fischer et al., 2013; Pannasch, Helmert, Roth, Herbold, & Walter, 2008; Unema, Pannasch, Joos, & Velichkovsky, 2005). This corresponds to well-known results of early holistic and subsequent analytic perception (Fitousi, 2020; Navon, 1977; Van Leeuwen & Bakker, 1995).

In our contour integration task, precursor fixations followed the largest saccades compared to other fixation categories. These large saccades may serve to mark locations in the visual stimulus that should be remembered for future revisits. These locations may either be highly salient or be difficult to acquire visual information from (Zhang et al., 2022), or they may contain an abundance of strategic information and therefore give rise to a plan to return later. Accordingly, Wilming and colleagues (2013) found precursor fixation locations to be more visually salient and fixated longer than ones the eyes do not return to. However, in our study fixation durations did not differ between fixation categories. This observation is inconsistent with both Wilming’s hypothesis that the visual system needs more time to pick up the salient information from these locations (Wilming et al., 2013), and Hooge’s hypothesis that the eye returns to precursor locations because of insufficient time for information uptake at the first visit (Hooge et al., 2005); in both these cases we would have expected precursor fixations to be shorter rather than longer. We might therefore consider the possibility that information uptake at precursor fixations differs qualitatively rather than quantitatively from other fixations (neither more nor less, but different). At the beginning of visual exploration, strategic locations may be spotted at the periphery and reached by large saccades. To build the exploration plan, information encoding during precursor fixations may operate in an exploratory, as opposed to an exploitative mode. Thus, precursor fixations, whose locations are encoded to preserve nodes of the visual scene for further visual exploration, may be considered pivotal moments in ongoing viewing.

The size of the saccades outgoing from the precursor fixations is much smaller than that of the incoming saccades (and is the smallest among the fixation categories) (Fig. 2C). This suggests that between these saccades, a shift may have occurred from an exploratory, holistic mode to an exploitative, analytic mode of perception. Once in this mode, fixation duration is now short (Fig. 2A) and saccade size is small, meaning that the region around the precursor fixation becomes subject to extensive scrutiny, consistently with Wilming and colleagues (2013).

It should be noted that the refixation behavior discussed above does not take into account immediate refixations or ‘lag-2 revisits’ (Godwin, Hout, Alexdottir, Walenchok, & Barnhart, 2021; Peterson et al., 2001; Zhang et al., 2022). These types of refixations occur right after the gaze leaves the object and are common in visual search. These immediate gaze returns act as corrective saccades when a fixation on the object was terminated before it was fully processed (Godwin et al., 2021). In our analysis, we used a criterion of 2° of visual angle radius to determine refixations (Fig. 1). Given the small distances between the Gabor patches, this inclusion criterion most likely filtered out most of the precursor fixations with immediate refixations, leaving only those that participated in the global exploration plan.

In sum, during lasting visual behavior, exploratory modes involve continuous creation and updating of the viewing plan, as evidenced by large saccades before and small saccades after the precursor fixations. Exploratory modes alternate with exploitative modes of scrutiny, when the gaze revisits previous locations. The repetitive transitions between these modes provide flexibility and adaptability to natural viewing.

The three eye movement categories studied are typical of unrestricted viewing behavior, which is characterized by repetitive gaze returns. The importance of refixations for goal-directed visual exploration is nowadays increasingly appreciated. This is supported by experimental studies that show the critical role of refixations in memory formation and retrieval in natural viewing (Kragel & Voss, 2022), as well as by major modeling projects that aim to simulate refixation sequences similar to those observed in multiple human and animal gaze datasets (Zhang et al., 2022). Our contribution to this research is limited to one particular task: contour integration in dense fields of equidistant and similar Gabor elements. It remains to be seen how our results generalize to different perceptual tasks. We have not attempted to relate precursor fixations to the peculiarities of the current task and to the properties of the visual scene and, whose competing influences are known to control eye movements (Tatler, Hayhoe, Land, & Ballard, 2011). Consequently, future research should explore the neural correlates of precursor fixations in natural scenes containing a broad spectrum of spatial frequencies and visual saliences and should be designed to analyze in more detail the brain processes around precursor fixations. Such research will have to take into account the overlapping effects that sequential eye movements produce on brain activity during unrestricted viewing. Our study shows that deconvolution modeling offers a robust solution to this problem. This can be inferred, in particular, from the successful elimination of the FRP difference between ordinary fixations and refixations related to the fixation rank, as well as from the elimination of the saccade size effect on the lambda peak, which is known to be sensitive to differences in saccade size (Dimigen et al., 2009; Kazai & Yagi, 1999; Thickbroom et al., 1991), which exist between all our fixation categories (Fig. 2C).

## Conclusions

Eye movements do not simply supply our vision with information or passively reflect the outputs of visual processing, but are themselves an inextricable part of the attention and memory brain systems (Awh, Armstrong, & Moore, 2006; Gottlieb, 2012; Voss, Bridge, Cohen, & Walker, 2017). Our study extends this understanding by showing the difference between the three basic categories of fixations in naturalistic viewing behavior. Precursor fixations to locations that are subsequently refixated have distinct neural correlates compared to refixations and ordinary fixations. While the neural mechanisms of refixations have been investigated earlier (Kragel et al., 2021; Meghanathan et al., 2020; Nikolaev et al., 2018), as far as we know, the neural mechanisms of precursor fixations have not been studied before. Their peculiarity may arise from the combined contribution of cognitive factors responsible for making a strategic plan of visual exploration, and oculomotor factors that determine the size of saccades depending on their serial position (rank) in a trial. These findings emphasize the intertwined character of cognitive processing and oculomotor behavior.

## Acknowledgments

ARN and RNM were supported by an Odysseus grant (G.0003.12) from the Flemish Organization for Science (FWO) to Cees van Leeuwen. ARN was also supported by a grant from the Marcus and Amelia Wallenberg Foundation (MAW2015.0043) to Mikael Johansson. BE was funded by Deutsche Forschungsgemeinschaft (DFG, German Research Foundation) under Germanýs Excellence Strategy – EXC 2075 – 390740016. We thank Nathalie Van Humbeeck and Johan Wagemans for permission to use the jointly collected data in this study. We thank Olaf Dimigen for his help with OPTICAT and in synchronizing EEG and eye movement recordings with the EYE-EEG extension for EEGLAB.

