## Supplementary material for "Planning to revisit: neural activity in refixation precursors"

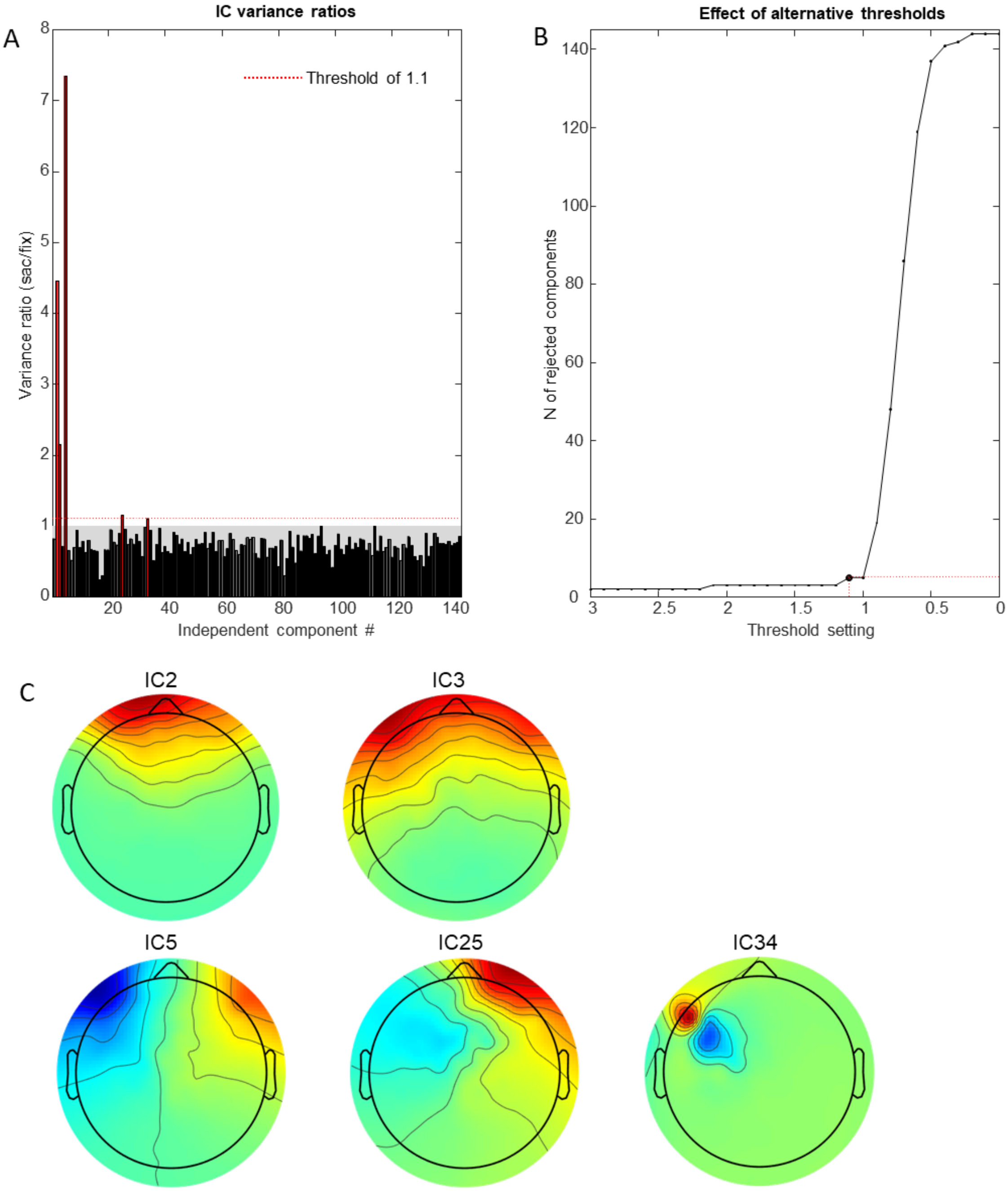


Fig. S1. Results of ocular artifact correction with the OPTICAT function (Dimigen, 2020) for a representative participant. A: Distribution of the variance ratio in saccade and fixation intervals of all independent components. If the ratio exceeded the threshold of 1.1, the component most likely reflected ocular artifacts and was rejected. Five such components were detected for this participant; they are marked red. B: Dependence of the number of rejected components on the threshold settings for this participant. The intersection point corresponds to the threshold 1.1 and to the five rejected components. C: Topographical maps of these five components, showing the predominantly frontal location of their maxima.

Table S1. The distribution of removed independent components by artifact category after the ICLabel classification per participant.

| | Mean | Min | Max | SD |
|---|---|---|---|---|
| Muscle, 0.4-1 | 5.2 | 1 | 13 | 3.4 |
| Eye, 0.9-1 | 0.1 | 0 | 1 | 0.4 |
| Heart, 0.05-1 | 2.5 | 0 | 6 | 1.8 |
| Line Noise, 0.4-1 | 0.9 | 0 | 6 | 1.4 |
| Channel Noise, 0.4-1 | 27.3 | 9 | 44 | 10.3 |
| Other, 0.4-1 | 51.7 | 26 | 85 | 14.6 |

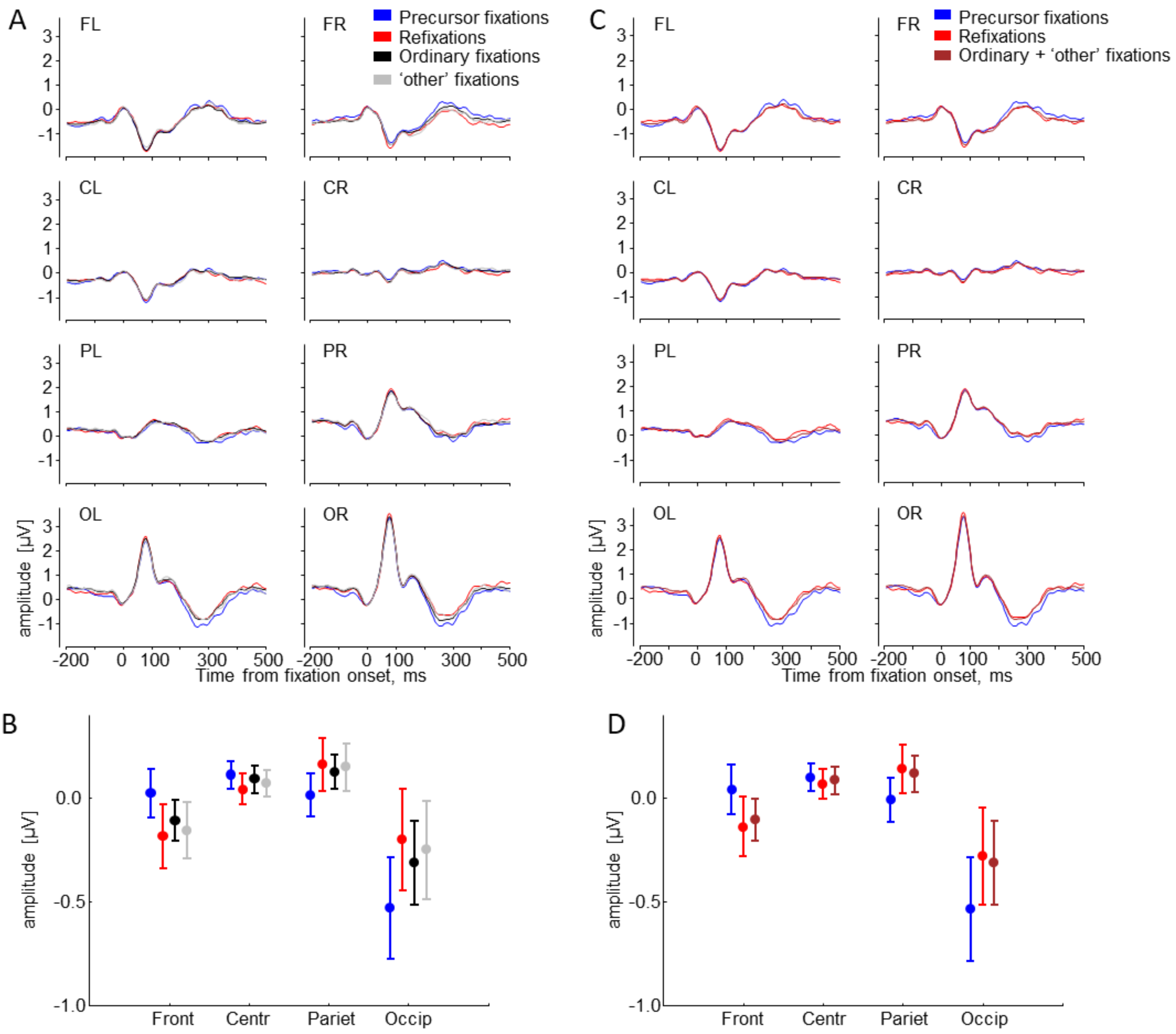


Fig. S2. The FRP results of the two additional analyses. A: Fixation-related potentials for 8 ROIs for four fixation categories, including the 'other' fixation category. B: Mean amplitude in the interval 200-400 ms after fixation onset for the four fixation categories (in A), indicating an interaction between fixation category and ROI. C: Fixation-related potentials for 8 ROIs for three fixation categories, where ordinary and 'other' fixations were combined before deconvolution. D: Mean amplitude in the interval 200-400 ms after fixation onset for the three fixation categories (in C), indicating an interaction between fixation category and ROI. The potentials are grand-averaged (N=21) and baseline-corrected at 0-20 ms after fixation onset. Error bars indicate standard errors of the means across 21 participants.
